# Engineered expression of the invertebrate-specific scorpion toxin AaHIT reduces adult longevity and female fecundity in the diamondback moth *Plutella xylostella*

**DOI:** 10.1101/2020.09.10.291187

**Authors:** T.D. Harvey-Samuel, X. Xu, E. Lovett, T. Dafa’alla, A. Walker, V.C. Norman, R. Carter, J. Teal, L. Akilan, P.T. Leftwich, L. Alphey

**Author notes:** **Authors for correspondence** Tim Harvey-Samuel -, Luke Alphey.

## Abstract

**BACKGROUND:** Previous Genetic Pest Management (GPM) systems in diamondback moth (DBM) have relied on expressing lethal proteins (‘effectors’) that are ‘cell-autonomous’ i.e. do not leave the cell they are expressed in. To increase the flexibility of future GPM systems in DBM, we aimed to assess the use of a non cell-autonomous, invertebrate-specific, neurotoxic effector – the scorpion toxin AaHIT. This AaHIT effector was designed to be secreted by expressing cells, potentially leading to effects on distant cells, specifically neuromuscular junctions.

**RESULTS:** Expression of AaHIT caused a ‘shaking/quivering’ phenotype which could be repressed by provision of an antidote (tetracycline); a phenotype consistent with the AaHIT mode-of-action. This effect was more pronounced when AaHIT expression was driven by the *Hr5/ie1* promoter (82.44% of males, 65.14% of females) rather than *Op/ie2 (*57.35% of males, 48.39% of females). Contrary to expectations, the shaking phenotype and observed fitness costs were limited to adults where they caused severe reductions in mean longevity (–81%) and median female fecundity (–93%). qPCR of AaHIT expression patterns and analysis of *piggyBac*-mediated transgene insertion sites suggest that restriction of observed effects to the adult stages may be due to influence of local genomic environment on the tetO-AaHIT transgene.

**CONCLUSION:** We have demonstrated the feasibility of using non cell-autonomous effectors within a GPM context for the first time in the Lepidoptera, one of the most economically damaging orders of insects. These findings provide a framework for extending this system to other pest Lepidoptera and to other secreted effectors.

## 1. Introduction

The diamondback moth - DBM (*Plutella xylostella*) is a highly invasive and economically important pest of Brassicas, costing upwards of $5 billion in damage and control measures each year (1, 2). Contributing to the notoriety of this pest is its extreme insecticide resistance with populations having developed resistance to most classes of insecticides, including *Bacillus thuringiensis* (*Bt*) Cry toxins and DDT (3). As such, research into novel, more environmentally friendly control methods is required.

Advances in molecular biology have allowed DBM to function as a model for the development of Genetic Pest Management (GPM) tools in the Lepidoptera. Previously, we developed a transgenic, tetracycline-repressible, female-specific lethality system in DBM based on the sex-specific alternative splicing circuitry of the *Pectinophora gossypiella* (pink bollworm) *doublesex* gene (4). This female-lethal system is repressed in the lab or rearing facility when larvae are fed a diet containing sufficient quantities of an antidote (tetracycline or suitable analogues – see Figure 1). In the generation prior to release, tetracycline is withdrawn from the diet resulting in the death of female larvae, allowing the release of adult males homozygous for the female lethal transgene which they then pass on to their progeny in the field after mating with wild females (5, 6). As tetracycline is not present in sufficient quantities under field conditions, female progeny inheriting the transgene will die prior to adulthood while male heterozygotes will survive to pass the transgene on to subsequent generations (7). Glasshouse trials have shown that repeated releases of these homozygous males can result in eradication of caged wild-type populations and, additionally, delay evolution of *Bt* resistance (8, 9), a finding in agreement with previous modelling (10, 11). Wind tunnel experiments have demonstrated that these males retain the ability to locate and respond to female pheromone plumes (12) and open-field trials have shown that these males are able to disperse within a realistic crop setting (13).

**Figure 1:**
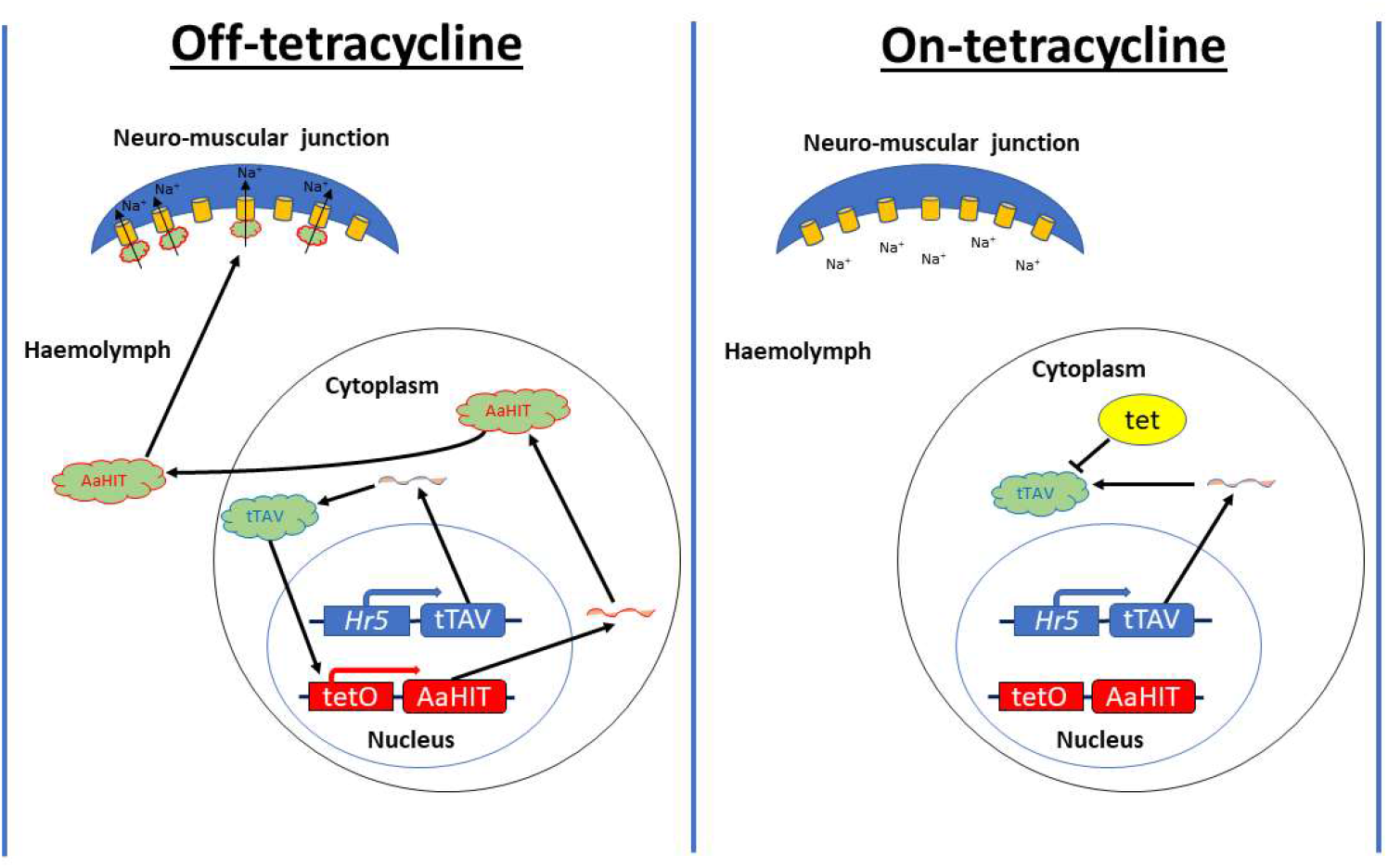
Schematic representation of the bipartite ‘tet-off’ system and its use in secreted neurotoxic effector (AaHIT) expression. This system is composed of two components 1) a transgene expressing tTAV (here represented by *Hr5*-tTAV) and 2) a transgene which in the presence of active tTAV will express the neurotoxic effector AaHIT (here represented by tetO-AaHIT). Left panel ‘Off tetracycline’: the system in its unrepressed state. In the absence of the antidote (tetracycline, tet), tTAV protein expressed from the *Hr5*-tTAV transgene will bind to tetO and drive the expression of AaHIT. The secretory peptide encoded within AaHIT allows it to leave the cell it is expressed in and travel through the haemolymph where it will encounter, bind and overstimulate neuromuscular junctions resulting in a loss of muscular control. Right panel ‘On-tetracycline’: This panel depicts the system in its repressed state. Here, tet is provided to the organism as it is developing. Tet binds the tTAV protein and prevents it binding to tetO thus inhibiting the expression of AaHIT and allowing individuals bearing this system to retain muscular control. In the context of using these systems as population control tools, the ‘On-tetracycline’ situation describes how strains bearing these transgenes would be mass-reared in the laboratory/rearing facility. The ‘Off-tetracycline’ situation describes how the progeny of released individuals developing in the field would be subject to fitness costs brought about through loss of muscle control.

Despite the rapid advance of this technology, a limitation – shared with a similar field-tested dominant lethal genetic system in the dengue fever mosquito *Aedes aegypti* (14) – is that it is based on the expression of ‘cell-autonomous’ effectors, for example the transcriptional activator tTAV from the ‘tet-off’ expression system (15). In an unrepressed state (e.g. in transgenic DBM female larvae in the field), expression of tTAV results in an uncontrolled positive feedback loop causing wide-scale gene misexpression/de-regulation, a situation which severely restricts the ability of the cell to perform its normal functioning (16). However, as the tTAV protein does not leave the cell in which it is expressed, the deleterious effects of its expression are limited to the cells in which its expression is directed (i.e. it is cell-autonomous), usually necessitating a design with a broad spatial expression range in order to ensure that this results in a sufficiently deleterious phenotype at the whole-organism level. This reliance on cell-autonomous effectors including tTAV has restricted the utility of more complex and intricate GPM systems utilising transcriptional regulatory elements (i.e. promoters and enhancers) which display useful temporal or spatially explicit expression patterns, for example if expression of tTAV in these areas would not necessarily result in organism death/non-viability (17-19).

In order to circumvent this limitation, we recently demonstrated a novel mechanism for the expression of non cell-autonomous effectors in the mosquito *Ae. aegypti* (20). There, fusion of the invertebrate-specific neurotoxic protein AaHIT – a component of the venom from the scorpion *Androctonus australis hector* – and the gp67 secretory signal peptide from the *Autographa californica* baculovirus resulted in a synthetic gp67-AaHIT effector protein (henceforth AaHIT) which could be secreted from cells it was expressed in and subsequently bind to voltage-gated sodium channels (VGSCs) at neuromuscular junctions, resulting in rapid onset of paralysis after expression (see Figure 1 for example of mechanism). The advantage of this system over those based on traditional cell-autonomous components is that as the effector is secreted, its ultimate effect (overstimulation of VGSCs) was independent of its original expression location (in the mosquito example, the cells of the adult female fat-body immediately post blood-feeding). This allowed the use of the female fat-body specific promoter from the *VitellogeninA1* gene even though expression of cell-autonomous effectors (Michelob, Reaper^KR^) using this promoter had no observable effect. The *AaHIT* gene has also been used to increase the potency of a variety of biocontrol technologies targeting lepidopteran pests including engineering its expression in recombinant *Bt* and *Autographa californica* baculovirus, both tested against DBM (21) and thus we hypothesised that such an AaHIT-based non cell-autonomous system could also function in this species. Here we extend our previous work in mosquitos to DBM using the same bipartite ‘tet-off’ gene expression system utilised in the DBM female-lethal GPM system. Utilising two transcriptional regulatory elements of known function in DBM (*Hr5/ie1* and *Op/ie2*) (22) to drive strong and ubiquitous tTAV production (henceforth *Hr5/ie1*-tTAV and *Op/ie2*-tTAV) we directed expression of a tetO*-AaHIT* transgene in DBM resulting in a tetracycline-repressible ‘shaking’ phenotype in adults that negatively affected their longevity and fecundity. These results lay the foundation for further research into a new generation of non cell-autonomous effector-based GPM strategies in this global pest and other lepidopterans.

## 2. Methods

### 2.1 Insect rearing

DBM larvae were reared on beet armyworm artificial diet (Frontier Biosciences, Germantown, Maryland, USA) under a 16 : 8 h light : dark cycle, 25° C and 50% relative humidity. Permissive conditions (henceforth on-tet) where the tet-off system is repressed were created by the addition of tetracycline-hydrochloride (Sigma, St Louis, Missouri, USA) to this diet to a final concentration of 1 μg/ml. Restrictive conditions (henceforth off-tet) where the tet-off system is unrepressed were created by making diet without tetracycline-hydrochloride. Adults stages were supplied with 10% sugar water *ad libitum* through soaked cotton wool. For on-tet conditions tetracycline-hydrochloride was added to this sugar water to a final concentration of 1 μg/ml.

### 2.2 Details of lines

Plasmids were generated using standard molecular biology techniques. Sequences for the three plasmids used (*Hr5/ie1*-tTAV (OX5378), *Op/ie2*-tTAV (OX4585) and tetO-*AaHIT* (AGG1074) were deposited into GenBank (Accession numbers MT533615, MT533614, and MT533613, respectively) and simplified schematics are given in Figure 2. Transgenic DBM lines were generated for each construct, and insertion copy number assessed, using previously published methods (22). Lines showing strong marker fluorescence were maintained as heterozygotes and a randomly selected line for each construct utilised in subsequent experiments. Genomic sequence flanking the *piggyBac* insertion site for each of the lines used here were identified using previously published methods (20, 22) (See Table 1) and the online NCBI BLAST programme NCBI https://blast.ncbi.nlm.nih.gov/Blast.cgi)

**Table 1:**
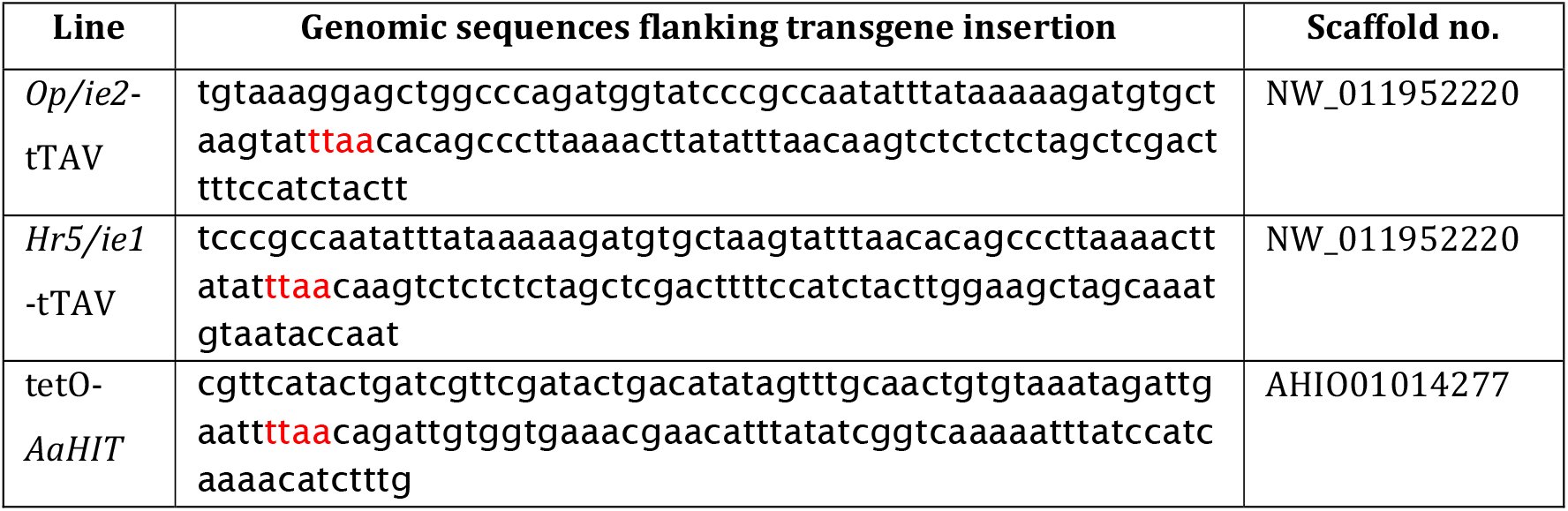
Genomic sequence flanking 60bp either side of TTAA insertion site (shown in red) for each of the lines used here.

**Figure 2:**
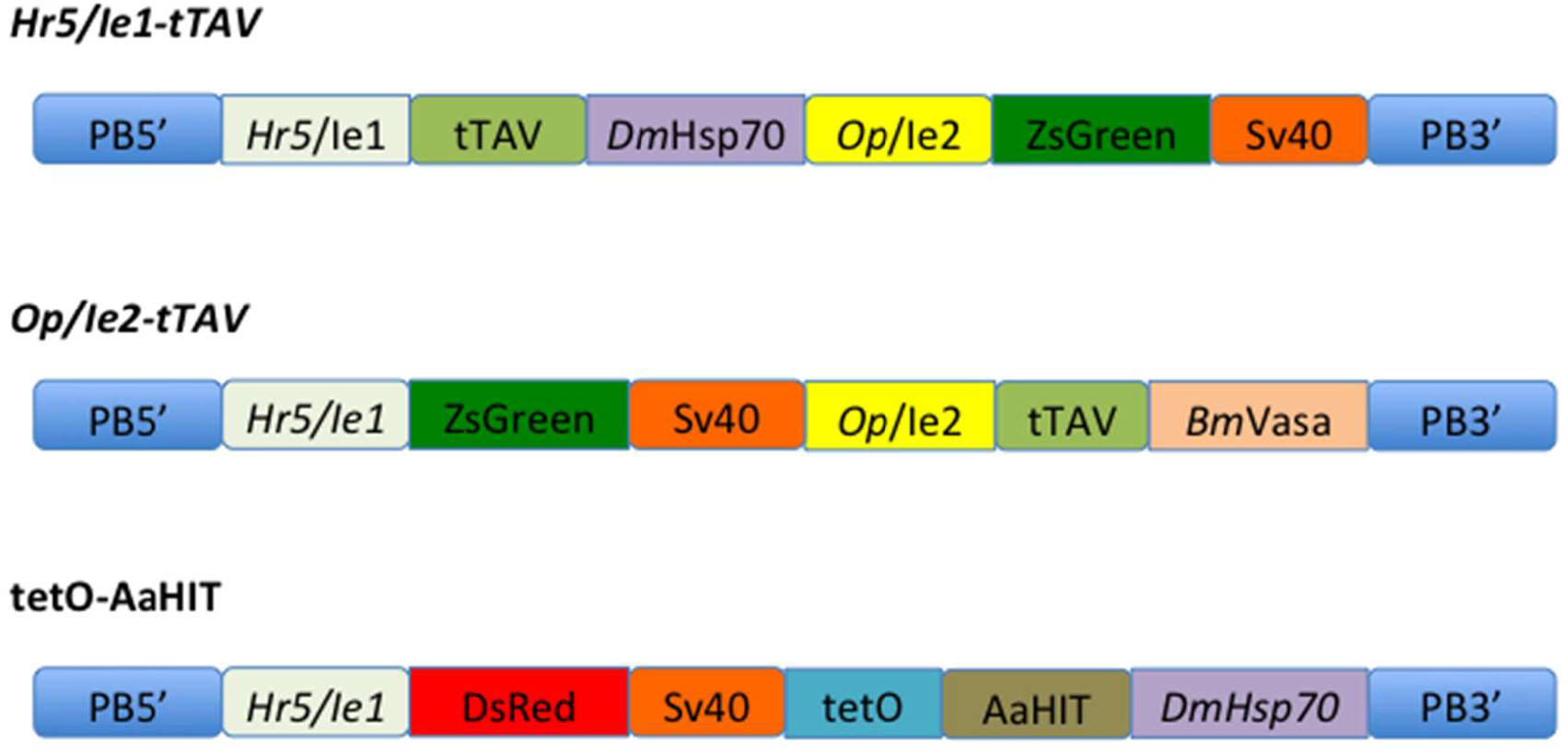
Simplified schematic representations of the three constructs used. Components are as follows: PB5’/PB3’ = *piggyBac* transposable element ITRs. *Hr5*/Ie1 = Homologous region 5 enhancer/immediate early gene 1 promoter+5’UTR from *A. californica* baculovirus. tTAV = tetracycline-controlled transactivator. *DmHsp70* = 3’UTR from *D. melanogaster Hsp70* gene. *Op/ie2* = Immediate early gene 2 promoter+5’UTR from *Orgyia pseudotsugata* baculovirus. ZsGreen = green fluorescent protein central transformation marker. Sv40 = 3’UTR from Simian virus 40. *BmVasa* = 3’UTR from *Bombyx mori vasa* gene. DsRed = red fluorescent protein central transformation marker. tetO = 7 repeats of the Tet operator sequence. AaHIT = gp67 secretory signal peptide from the *A. californica* baculovirus fused to *A. hector* neurotoxic venom protein.

**Figure 3:**
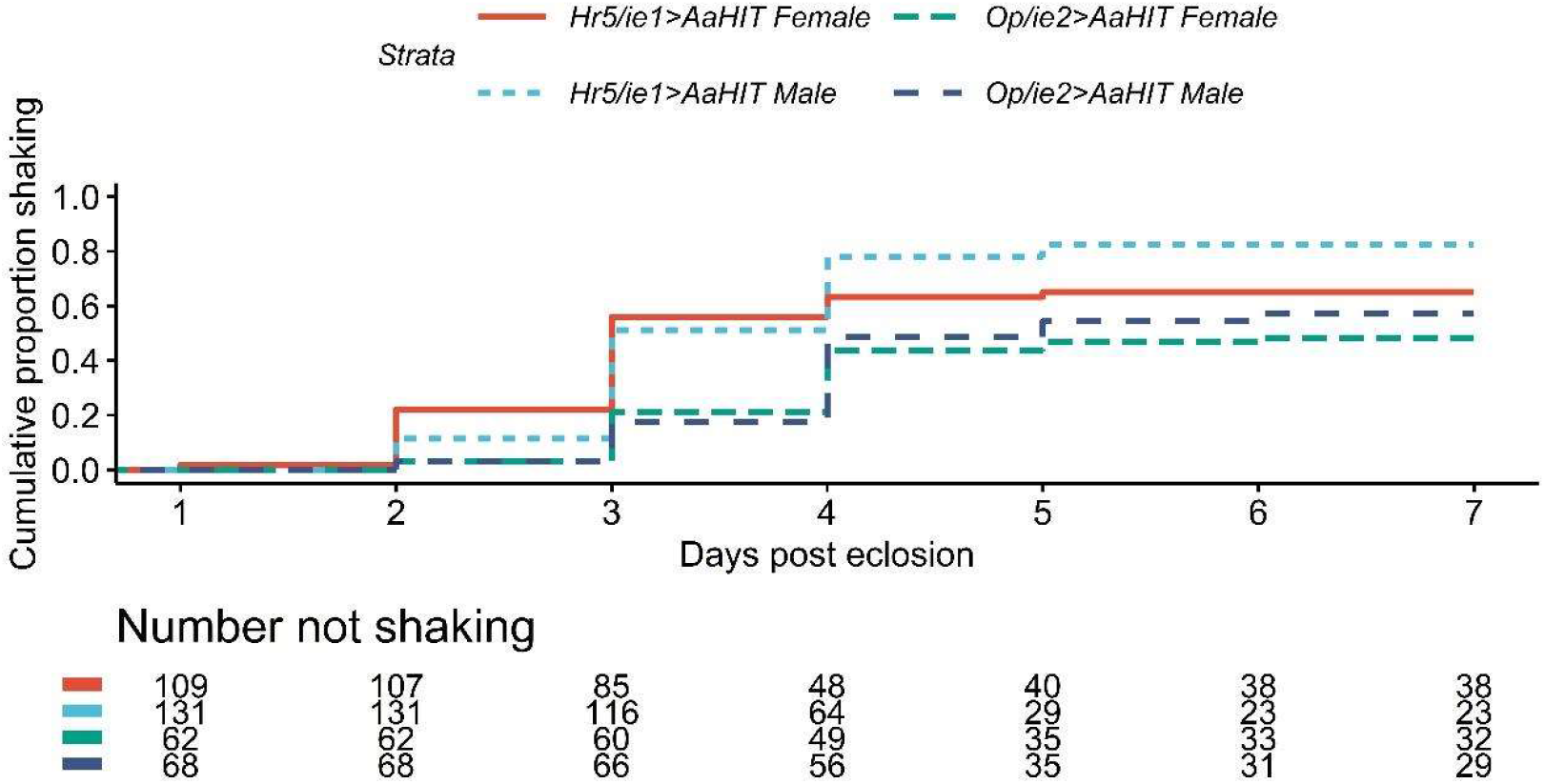
Expression of AaHIT causes an adult ‘shaking’ phenotype. Upper panel: Cumulative proportion of adults in each cohort recorded as shaking each day over the seven day observation period. This figure only displays results for the four double heterozygous cohorts (i.e. *Hr5/ie1*>*AaHIT* and *Op/ie2*>*AaHIT* males and females) as no shaking events were observed in any of the other 28 cohorts assessed. Lower panel: data from which cumulative proportions were calculated. Results analysed using a Cox’s Proportional Hazards test suggesting *Hr5/Ie1*>*AaHIT* individuals were significantly more likely to display a shaking phenotype than *Op/ie2*>*AaHIT* individuals (n =370, Z = −5.11, p<0.005) but no significant effect of sex.

### 2.3 Survival to eclosion experiment

Heterozygous tetO-*AaHIT* adult females and heterozygous *Hr5/ie1*-tTAV adult males were mated *en masse*, with the reciprocal cross also conducted in a separate cage. Adults were supplied with cabbage-juice painted Parafilm (Bemis Company Inc. Neenah, Wisconsin, USA) to act as an oviposition substrate. After 24 hrs, egg sheets were removed from each cage and divided into two pieces. One piece from each cross was then placed onto either on-tet or off-tet diet. On and off-tet cohorts were kept separate for the remainder of the experiment and analysed separately. At pupation all individuals were screened for fluorescence and subsequently separated into the four possible genotypes i.e. *Hr5/ie1*-tTAV single heterozygotes, *Hr5/ie1*-tTAV + tetO-*AaHIT* double heterozygotes (henceforth *Hr5/ie1*>*AaHIT*), tetO-*AaHIT* single heterozygotes and wild-type (WT). Cohorts were allowed to eclose and numbers of pupae and subsequent adults compared between genotypes based on an expected 1:1:1:1 ratio (Chi-square goodness of fit). The same experiment was performed using the *Op/ie2*-tTAV line in place of *Hr5/ie1*-tTAV.

### 2.4 Adult time series experiment (analysis of shaking phenotype)

Heterozygous *Hr5/ie1*-tTAV and heterozygous tetO-*AaHIT* individuals were crossed in the same way as above to give pupae of the four possible genotypes, reared on and off tetracycline and separated by sex (16 cohorts total). However, as pupae, each cohort was placed into a cage (BugDorm, Taiwan) with added sugar water soaked cotton wool of the appropriate tetracycline condition. These pupae were observed daily for eclosion with the first day of adult eclosion counted as Day 0 of the experimental period for that cage – analysis was conducted from the first day post eclosion (1 dpe). As cohorts were separated by sex, eclosion timing was relatively simultaneous in each cage. From this period, all cages were observed daily and any adults which displayed a shaking phenotype were removed from the cage, placed in RNA*later* (Invitrogen, Carlsbad, California, USA) and frozen in liquid nitrogen for subsequent analysis. Observations continued until all individuals were dead or collected in the *Hr5/ie1*>*AaHIT* off-tet cage. The same experiment was also performed using the *Op/ie2*-tTAV line in place of *Hr5/ie1*-tTAV. Individuals which failed to eclose were removed from each cohort dataset prior to analysis. Analysis of cohorts which had shown a phenotype was conducted using a Cox’s Proportional Hazards test comparing between tTAV lines and sexes.

### 2.5 Adult longevity experiment

This experiment was conducted using the *Hr5/ie1*-tTAV and tetO-*AaHIT* lines. The 16 cohorts were created as described above. Thirty pupae from each of the *Hr5/ie1*>*AaHIT* on- and off-tet, male and female cohorts were placed in individual observation pots, supplied with sugar water of the appropriate tetracycline condition and observed daily. Recorded for each individual was the day of eclosion and the day of death. From this data a longevity analysis was performed using a Cox’s Proportional Hazards test.

### 2.6 Female fecundity experiment

This experiment was conducted using the *Hr5/ie1*-tTAV and tetO-*AaHIT* lines. The 16 cohorts were created as described above. Thirty pupae of each of the *Hr5/ie1*>*AaHIT* on- and off-tet female cohorts were placed in individual observation pots, supplied with sugar water of the appropriate tetracycline condition and observed daily. On the day each female eclosed, three freshly eclosed, virgin WT males were placed in each individual pot. Females were allowed to mate and lay eggs on cabbage-juice painted parafilm for 72 hrs after which time egg sheets were collected. Eggs were counted on each sheet and hatch rates calculated. Statistical differences in egg laying and hatch rate were assessed using a quasipoisson and quasibinomial glm, respectively.

### 2.7 qPCR experiment

#### 2.7.1 Total RNA extraction and cDNA synthesis

Total RNA from individual whole DBM from various life stages (L2 off-tet *n*=3, L2 on-tet n=3, L4 off-tet *n*=3, L4 on-tet n=3, Pupae off-tet n=3, Pupae on-tet n=3, 2 dpe off-tet n=4, 2 dpe on-tet n=4, 3 dpe off-tet n=4, 3 dpe on-tet n=3, 4 dpe off-tet n=2, 4 dpe on-tet n=3) was extracted using the Qiagen RNeasy Mini Plus Kit (Qiagen, Hilden, Germany), which includes a column-based genomic DNA removal step. Extracted RNA was quantified using a NanoDrop 2000/2000c Spectrophotometer (ThermoFisher Scientific, Waltham, Mass., USA). Complementary DNA (cDNA) was synthesized from total RNA using the High Capacity cDNA Reverse Transcription Kit (Applied Biosystems, Foster City, CA, USA) following the manufacturers protocol. After reverse transcription cDNA was stored at −20 °C.

#### 2.7.2 RT-qPCR assay design

Primers for 40S ribosomal protein S17 gene (*17S;* NM_001305512.1) were manually designed after interrogation of the genomic sequence (downloaded from http://iae.fafu.edu.cn/DBM/). Primer sets for elongation factor 1 gene (*EF1*; XM_011562844.1), ribosomal protein L32 gene (*RpL32*; NM_001309136.1), and ribosomal protein S13 gene (*RpS13*; NM_001305523.1) were previously designed by (23). Primers for *AaHIT* were manually designed in-house. Summarised in Table 2.

Each RT-qPCR run was setup manually by the same operator in MicroAmp fast optical 96-well reaction plates with barcode 0.1 mL (Applied Biosystems) and sealed with MicroAmp optical adhesive film (Applied Biosystems) at The Pirbright Institute and run on the QuantStudio3 PCR system (Life Technologies, Carlsbad, CA, USA) on ‘Fast’ cycling conditions using the QuantStudio Design and Analysis Software 1.3.1. Each reaction was run in 10μL volumes and included 1x Luna Universal qPCR Master Mix (New England BioLabs, Ipswich, Mass. USA), optimal forward and reverse concentrations were experimentally determined (200nM for both forward and reverse primers for *17S, EF1, RpS13*, and *AaHIT*; 150nM for both forward and reverse primers for *RpL32*), and 1μg of template. Reactions were run in triplicate, including no template and no-RT controls. The cycling conditions were as follows: an initial 95°C for 1 min, followed by 95°C for 5 sec, and 60°C for 20 sec for 40 cycles.

#### 2.7.3 RT-qPCR - analysis

Standard curves, including the slope, y-intercept, correlation coefficient, and the PCR efficiency for each primer set was constructed using the QuantStudio Design and Analysis Software 1.3.1.

A geNORM analysis was conducted to determine which reference genes out of the four measured (*17S, EF1, RPL32, RPS13*) were the most stably expressed across all life stages and treatment groups. Raw RT-qPCR data was read into the statistical computing software R Version 1.2.5033 using the package “ReadqPCR” (Perkins et al. 2012), and the geNORM analysis was conducted using the package “NormqPCR” (24). The four reference genes were ranked across all life stage by treatment group combinations based on their gene stability measure (M), and the gene with the highest M value was excluded. The average expression stability was then calculated for each life stage and treatment group combination, as was the pairwise variation. The three most stable (17S, RPL32, EF1), and therefore most suitable, reference genes were used to normalize the Cq values using the qBase relative quantification framework (25). NRQ values were calculated by making each normalised biological sample (mean of technical triplicates) relative to an independent calibrator consisting of equimolar pooled cDNA from all 38 samples, itself also a mean of technical triplicates. For comparisons between life stages on and off-tet (i.e. Figure 4), only sampling points where at least 3 biological replicates were present in both on and off-tet treatments were included (33 samples).

**Table 2.**
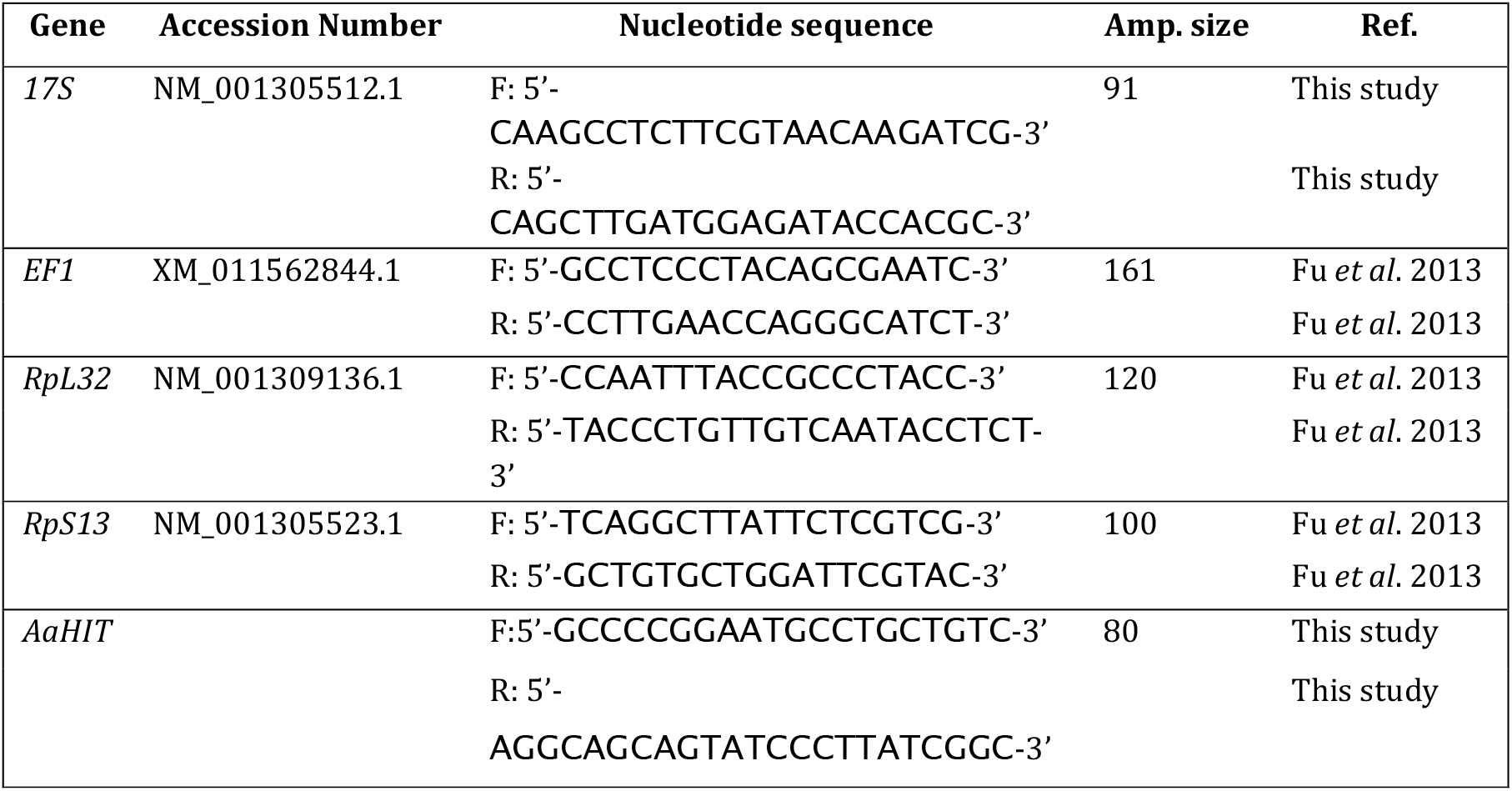
Description of RT-qPCR primer sequences, genomic location and amplicon characteristics.

**Figure 4.**
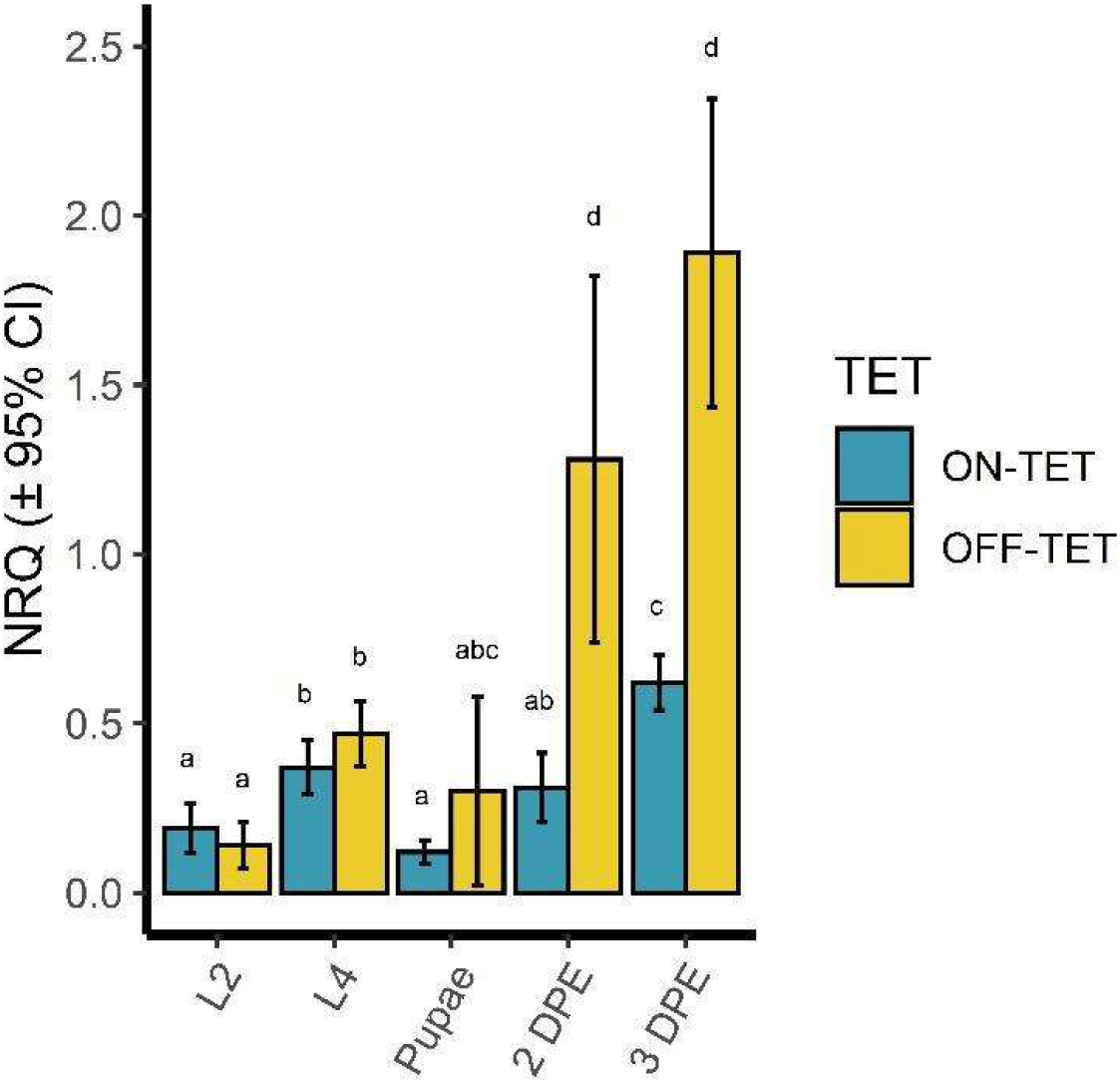
qPCR analysis of AaHIT expression. Normalized Relative quantification (NRQ) values of AaHIT expression from *Hr5/ie1*>*AaHIT* DBM reared on and off-tet across several life stages - L2 larvae, L4 larvae, pupae, 2 & 3-dpe (days post eclosion) adults. Bars represent the geometric means of biological replicates for relative gene expression. Error bars are 95% confidence intervals and lowercase letter groupings denote significantly different groups as denoted by non-overlapping confidence intervals. Sample sizes provided in methods.

### 2.8 Statistics

Data analysis was conducted with R v3.6.2 (R Core Team 2019). Survival analyses were conducted with the *survival* (26), *survminer* (27), packages. Figures were made with *ggplot2* from the tidyverse range of packages (29).

## 3. Results and Discussion

Previous studies involving the engineering of *AaHIT* into baculoviruses and *Bt* have demonstrated a significantly increased lethality to larval stages of DBM (21). As such we first set out to assess whether the transgenic expression of the synthetic AaHIT neurotoxin affected survival to pupation and/or adult eclosion. *Hr5/ie1*-tTAV and *Op/ie2*-tTAV heterozygotes were crossed to tetO-*AaHIT* line heterozygotes *en masse* (e.g. *Hr5/ie1*-tTAV males x tetO-*AaHIT* females and vice versa). Egg sheets from each cross were divided in half and reared either on-tet or off-tet. Pupae from each cross were collected, separated by genotype through screening for fluorescence markers, counted and then allowed to eclose. From each cross, given equal survival, pupae of four genotypes should have been present (e.g. *Hr5/ie1*-tTAV single heterozygotes, tetO-*AaHIT* single heterozygotes, *Hr5/ie1*>*AaHIT* double heterozygotes - and wild-type) at a 1:1:1:1 ratio. Deviation from this ratio indicates potential fitness costs in the under-represented genotypes. Overall, we found no significant deviation from this ratio for any of the treatment conditions (see Table 3). Similarly, the number of adults eclosing from these pupae did not significantly differ from the expected 1:1:1:1 ratio (see Table 4).

**Table 3:**
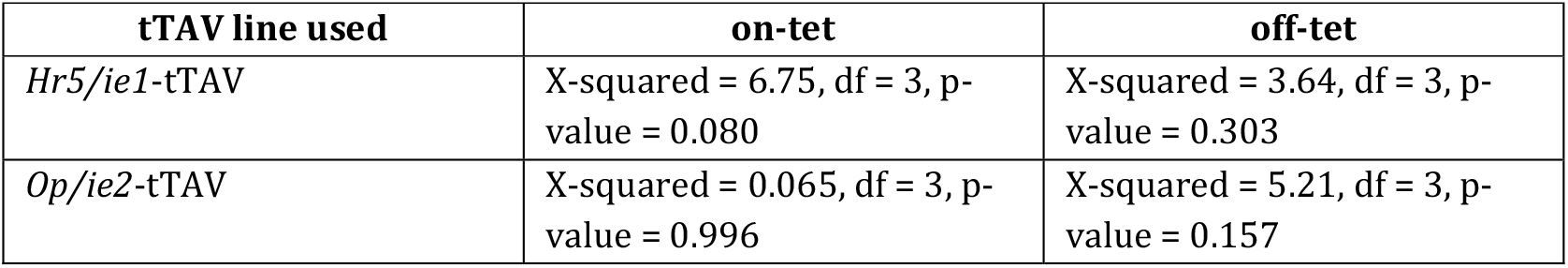
Results of Chi-square test of given probabilities comparing numbers of pupae recorded across the four genotypes and an expected 1:1:1:1 ratio. Each combination of tTAV line and tetracycline treatment analysed separately. No significant differences observed from expected ratio.

| tTAV line used | on-tet | off-tet |
| --- | --- | --- |
| *Hr5/ie1*-tTAV | X-squared = 6.75, df = 3, p-value = 0.080 | X-squared = 3.64, df = 3, p-value = 0.303 |
| *Op/ie2*-tTAV | X-squared = 0.065, df = 3, p-value = 0.996 | X-squared = 5.21, df = 3, p-value = 0.157 |

**Table 4:**
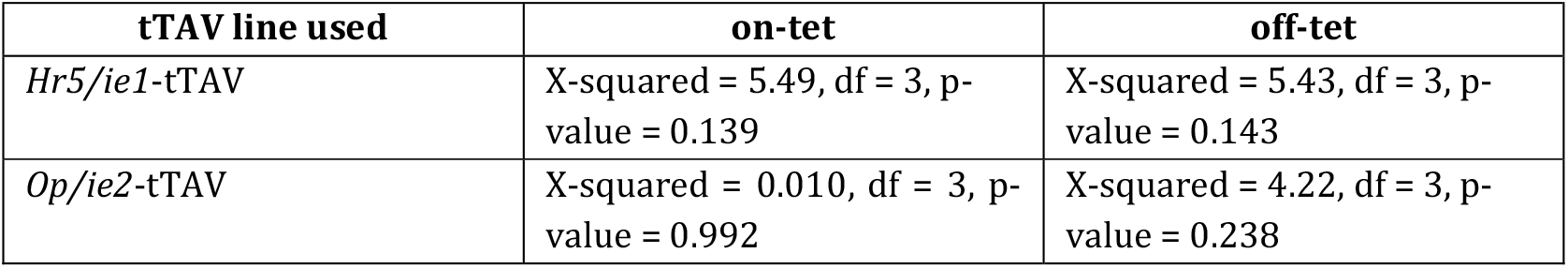
Results of Chi-square test of given probabilities comparing numbers of adults recorded across the four genotypes and an expected 1:1:1:1 ratio. Each combination of tTAV line and tetracycline treatment analysed separately. No significant differences observed from expected ratio.

| tTAV line used | on-tet | off-tet |
| --- | --- | --- |
| *Hr5/ie1*-tTAV | X-squared = 5.49, df = 3, p-value = 0.139 | X-squared = 5.43, df = 3, p-value = 0.143 |
| *Op/ie2*-tTAV | X-squared = 0.010, df = 3, p-value = 0.992 | X-squared = 4.22, df = 3, p-value = 0.238 |

With the absence of an observed effect on survival at larval/pupal stages, we next tested whether a phenotype consistent with the mode of action of AaHIT could be detected in adult stages. Reciprocal crosses were again performed as previously giving eight pupal genotype cohorts for each of the two tTAV line (e.g. for the *Hr5/ie1*-tTAV line - four genotypes on- and off-tet = eight cohorts). All pupae were subsequently sexed giving a total of 16 cohorts per tTAV line (i.e. 32 cohorts total). Pupae from each cohort were placed in a separate cage and observed daily. No abnormal phenotype was observed in any of the single heterozygote or wild-type cages of either sex or tetracycline treatment. However, beginning 1 day post eclosion (1 dpe) for the *Hr5/ie1*>*AaHIT* cohort and 2 dpe for the *Op/ie2*>*AaHIT* cohort, adults were observed displaying a ‘shaking’ phenotype typified by uncontrolled and rapid vibration of the wings and body and uncoordinated locomotion (see Figure 3. and videos 1-12 (Supporting Information)). Additionally, it was anecdotally observed that these cages had fewer flying individuals (i.e. more of their individuals were stationary on the floor/walls of the cage), although this observation was not assessed statistically. After being recorded as shaking and sexed by eye, adults displaying this phenotype were placed individually in RNA*later* (Invitrogen, Carlsbad, Ca, USA). and stored at −80° C for subsequent analysis. In each cohort, the number of individuals displaying this phenotype (per day) increased for two days at which point it reached its peak (*Hr5/ie1*>*AaHIT*, 3 dpe: 39.9% of males and 33.9% of females started shaking. *Op/ie2*>*AaHIT*, 4 dpe: 30.9% of males and 22.6% of females started shaking – in both cases percentages are of the starting number of individuals of that sex in that cage, not the number of remaining individuals. Over the seven days of observation a total of 82.4% of males and 65.1% of females from the *Hr5/ie1*>*AaHIT* cohort and 57.4% of males and 48.4% of females from the *Op/ie2*>*AaHIT* cohort were observed shaking. Observations ceased at 6 dpe and 7 dpe for *Hr5/ie1*>*AaHIT* and *Op/ie2*>*AaHIT*, respectively, as at this point all adults in these cages had either been recorded as shaking (and thus removed) or were dead. An analysis of the shaking behaviour observed (number of shaking events over time) using a Cox’s Proportional Hazards Test suggested a strongly significant effect of tTAV line on shaking hazard (*Hr5/ie1*>*AaHIT* cohorts 1.57-2.7 times (57% - 170%) more likely to display shaking behaviour than *Op/ie2*>*AaHIT* cohorts) (n =370, Z = −5.11, p<0.005) but no significant effect of sex on shaking hazard at this level of replication (n= 370, Z = 1.8, p = 0.073). From an applied standpoint, these results would suggest further work into non cell-autonomous expression systems in DBM may benefit from the prioritisation of the *Hr5/ie1* promoter over the *Op/ie2* promoter. However, to assess the generality of this conclusion further analysis of other insertion sites bearing these constructs to account for any potential positional effects would be required.

Given that the *Hr5/ie1* and *Op/ie2* promoters are known from previous work to express from early stages in DBM (e.g. from eggs), including as directors of fluorescent protein marker expression in the two lines used here, we were intrigued as to why the shaking phenotype observed was confined to adult stages. In order to explore this we performed qPCR analysis of AaHIT expression using a selection of those individuals that had been collected from the on and off-tet *Hr5/ie1*>*AaHIT* cohorts during the previous experiment (Figure 4).

These results aligned with the observed trends in shaking behaviour. In off-tet samples, levels of AaHIT transcript were relatively low at pre-adult stages before rising in the post eclosion (adult) samples. The highest levels of AaHIT expression coincided with the peak of the shaking phenotype observed in the previous experiment (3 dpe). In general, levels of *AaHIT* expression in on-tet samples were relatively low and did not overlap with levels observed in any off-tet shaking individuals. A caveat of these experiments is that while pre-adult samples were selected at random from individuals of the relevant genotype in the developing population, adult samples represent a random selection from individuals which were observed shaking on that day (in the off-tet groups). As such, the estimated NRQ values of these adult samples may represent the upper level of AaHIT expression amongst the *Hr5/ie1*>*AaHIT* adults as a whole.

These results suggest that the lack of observed shaking or other deleterious effects observed in pre-adults was possibly due to a lack of significant upregulation in AaHIT expression during these stages. An alternative explanation is insensitivity of the pre-adulthood stages to the AaHIT toxin, however this is deemed unlikely given the previously successful transgenic use of this toxin against early larval stages of DBM and other lepidopterans (21).

Normally, it would be expected that expression of a tetO-transgene would mirror the expression profile of the tTAV transgene(s) present, possibly with some additional basal expression of the tetO-transgene. The baculovirus promoters chosen here to regulate tTAV are known to display strong, constitutive and ubiquitous expression in DBM and indeed conformed to these expectations when driving the fluorescent marker (*ZsGreen*) in each tTAV line, which could be observed from late egg stages onwards (data not shown). It is therefore possible that positional effects caused by the tetO-*AaHIT* insertion site may have played a role in restricting substantial AaHIT expression to adults (e.g. through chromatin structure reducing access of tTAV or other transcriptional machinery to relevant areas prior to these stages). Characterisation of the tetO-*AaHIT* insertion site revealed it was located within the putative 5’ regulatory region of the *Aldehyde oxidase 1-like* gene (LOC105383938). The closest *B. mori* match to this gene - *xanthine dehydrogenase 1* (LOC101738209) shows substantial upregulation immediately prior to pupation. If this upregulation was related to a heterochromatin-euchromatin transition, and this situation was mimicked in DBM, this may possibly explain the adult-specific response of the tetO-AaHIT line observed here. Additional characterisation of other tetO and tTAV transgene insertions sites would be required to address this question further.

In addition to being of fundamental interest, the above results are consistent with previous findings in A*e. aegypti* in demonstrating the speed with which convulsion/paralysis phenotypes can be brought about using this synthetic non-cell autonomous system - in both cases, almost coincident with significant AaHIT upregulation. These findings bode well for the use of this system in situations where rapid-onset of toxicity in the pest species is required.

Although we did not observe a deleterious phenotype in larvae or pupae, we hypothesised that the rapid onset of the shaking phenotype in adult stages may negatively impact one or more fitness components. In order to explore this, we next assessed relative adult male and female longevity and female fecundity (number of eggs laid and hatch rate) in *Hr5/ie1*>*AaHIT* individuals.

In order to assess effects of AaHIT expression on adult longevity, on- and off-tet cohorts of *Hr5/ie1*>*AaHIT* pupae were first generated and sexed as previously described. A minimum of 24 pupae from each of the four cohorts (on and off-tet, male and female) were individually placed in observation pots and their dates of eclosion and death recorded. Using a Cox’s Proportional Hazards Test we found a highly significant effect of tetracycline provision on adult longevity (n = 102, Z = −6.00, p<0.005) with off-tet adults living an average of 3.23 ± 0.137 SE days and on-tet adults an average of 16.7 days ± 1.10 SE (reduction in average longevity of c. 81%) (see Figure 5). We observed no significant interaction between tetracycline provision and sex at this level of replication – marginal effect (n= 102, Z = −1.89, p= 0.059).

**Figure 5:**
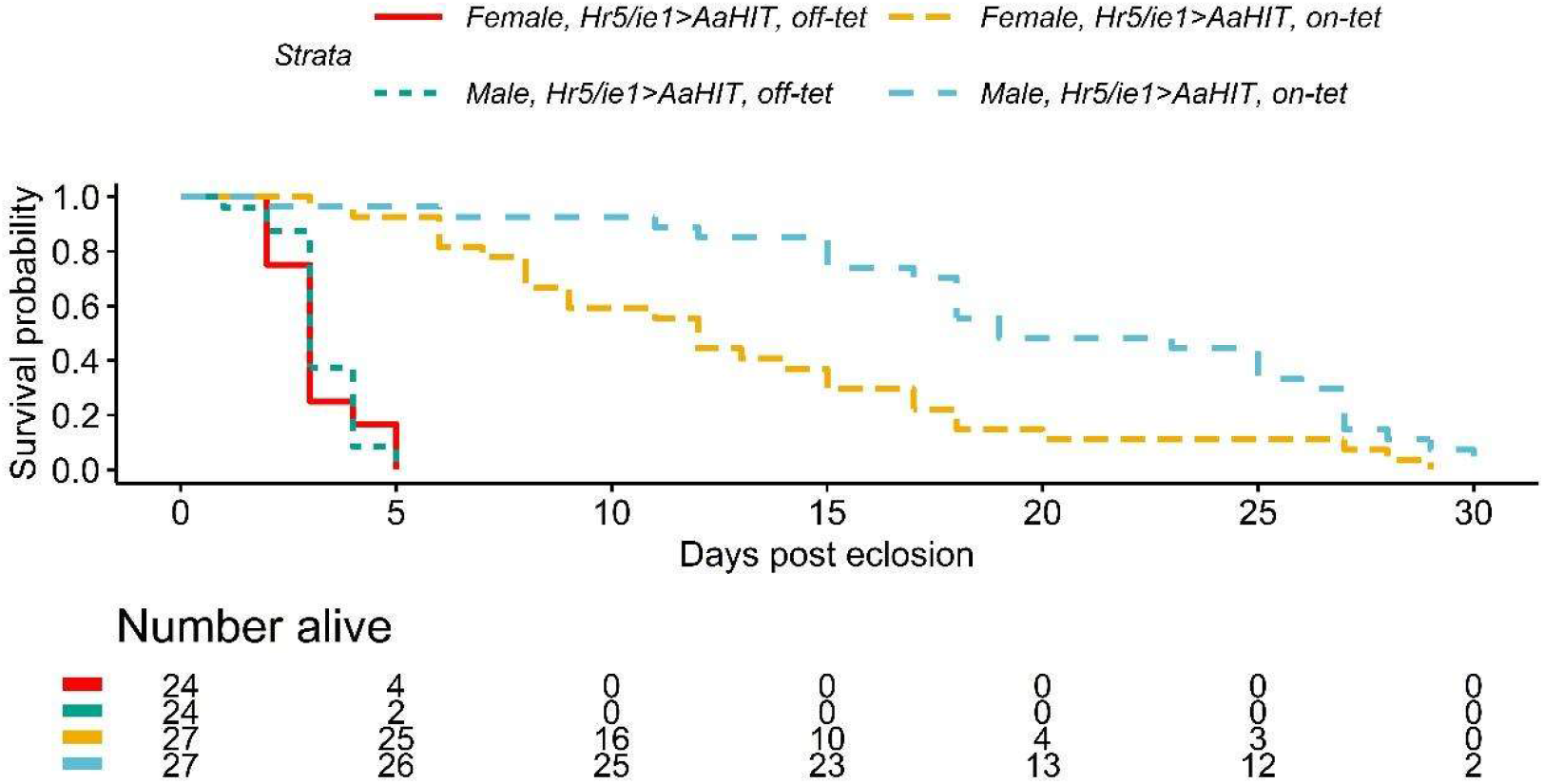
Effect on adult longevity of AaHIT expression. Upper panel: Daily survival probability for adults from four cohorts (*Hr5/ie1*>*AaHIT* male and females, on- and off-tet). Lower panel: data from which survival probabilities were calculated. Results analysed using a Cox’s Proportional Hazards test suggesting a significant effect of tetracycline treatment on adult longevity (n = 102, Z = −6.00, p<0.005) but no significant interaction between sex and tetracycline treatment at this level of replication.

In order to assess effects of AaHIT expression on female fecundity, sexed *Hr5/ie1*>*AaHIT* cohorts were first produced as above. A minimum of 26 individual female pupae from each tetracycline provision were placed in observation pots and on the day of eclosion, provided with 3 virgin, WT adult males. Females were allowed to mate and lay eggs for 72hrs (c. the average lifespan of off-tet individuals from the previous experiment) after which time laid eggs were removed and counted with hatch rates also recorded. Using a quasipoisson glm we found a significant effect of tetracycline provision on the number of eggs laid by females over this time period (n = 55, t = 8.06, p<0.005) with on-tet females laying a median of 124.5 eggs and off-tet females a median of 20 eggs (reduction in median eggs laying of c. 83.9%) see Figure 6. We further observed a significant effect of tetracycline provision on proportion of eggs hatched (n = 55, t = 4.05, p<0.005 – quasibinomial glm) with a median of 0.95 of eggs laid by on-tet females hatching but only 0.82 of eggs laid by off-tet females hatching (reduction in median proportion hatched of 13.7%). Overall, the median fecundity (number of larvae produced) of off-tet females (8.5) was reduced by c. 93% compared to on-tet females (119.5).

**Figure 6:**
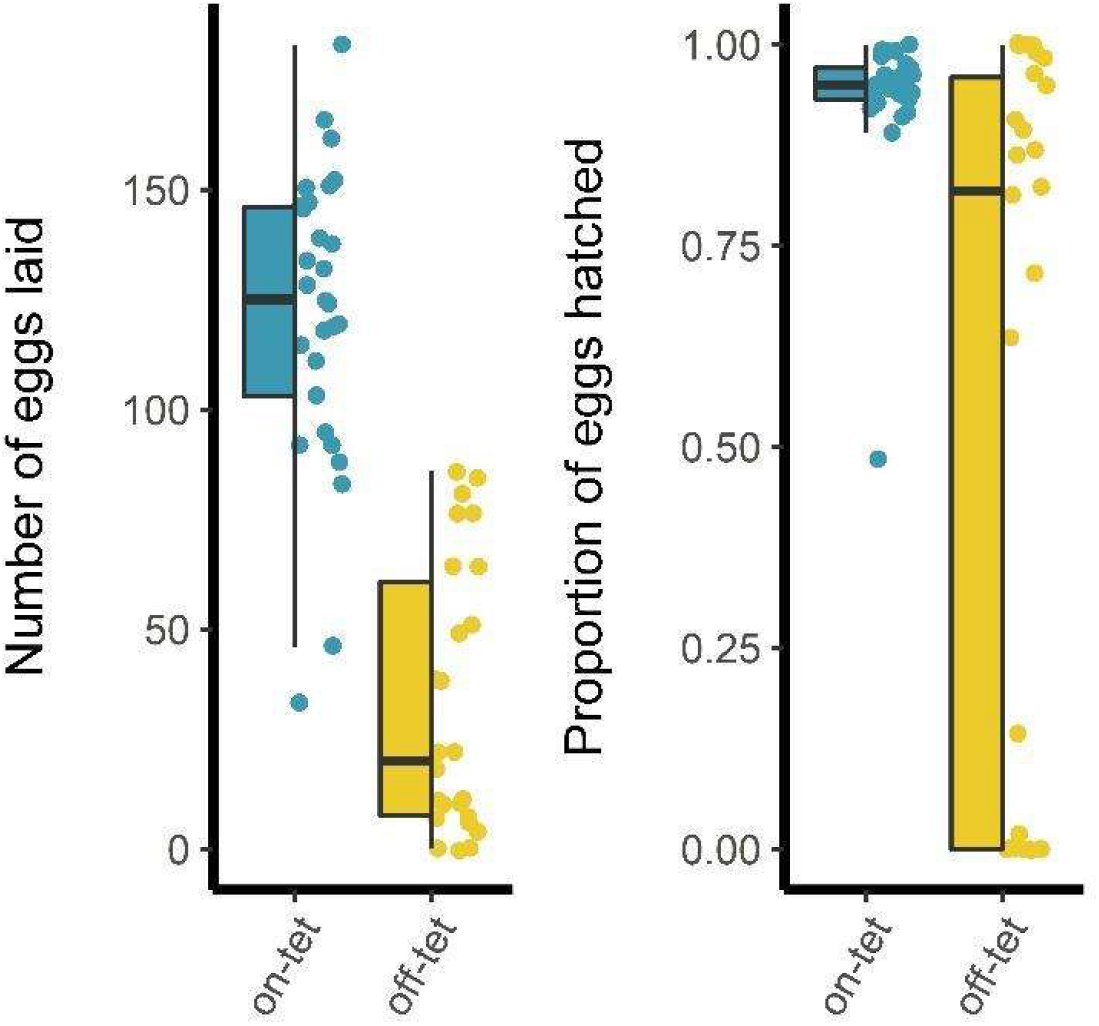
Effects on Female fecundity of AaHIT expression. Left panel: Number of eggs laid by *Hr5/ie1*>AaHIT females reared either on or off-tetracycline. Off-tet females laid significantly fewer eggs (median = 20 eggs) than on-tet females (median = 124.5 eggs), n = 55, Z = 7.54, p<0.005 – quasipoisson glm. Right panel: Of these eggs, a significantly lower proportion hatched when laid by off-tet females (median = 0.82) than on-tet females (median = 0.95), n = 55, Z = 4.39, p<0.005 – quasibinomial glm. Box limits represent the 25^th^ and 75^th^ percentiles, whiskers represent 1.5X the interquartile range, middle tendency estimates represent the medians.

These results contrast with those previously observed in *Ae. aegypti* where expression of AaHIT did not result in significant effects on adult female survival or egg-laying (hatch rates were not recorded in that experiment). A possible reason for this lies in the difference in how AaHIT was expressed in these two experiments. In *Ae. aegypti* we employed the *VitellogeninA1* promoter element to drive tTAV (and therefore AaHIT) production (30-32). This promoter shows a short window of upregulation (approx. 24hrs) in adult female fat-body cells immediately following a blood meal. On the other hand, here we used the less restricted baculovirus promoters *Hr5/ie1* and *Op/ie2*. Although the expected expression profiles of these two promoters did not drive a concomitant upregulation of AaHIT in stages prior to adults (possible reasons for this addressed above) it is possible that, once active, the tetO-insertion will have expressed in a larger number of cells and over a longer period than in our *Ae. aegypti* experiments. Therefore, where previously we achieved a short period of adult female paralysis followed by almost total recovery, here the induction of the tetO-AaHIT transgene resulted in reduced egg production and early death. This complex interplay between component choice and the confounding effects of transgene genomic context highlight both the flexibility possible in developing such GPM strategies but also the importance of individual component testing.

## 4. Conclusion

We have demonstrated the first use of a non cell-autonomous transgene-based effector in a lepidopteran, causing adult-specific, tetracycline-repressible, neurotoxic effects. Diamondback moth adults experiencing these neurotoxic effects were observed to be less mobile; often sitting on the bottom or wall of their enclosure with a shaking/shivering phenotype. On its own, it is possible that such a phenotype would impose significant fitness costs under real-world conditions where it could impede their ability to avoid an array of biotic and abiotic dangers. However, even in the relatively safe and stable environment of the laboratory, these adults experienced substantial decreases in their longevity and in female fecundity – both relevant parameters in terms of population control. These effects would, for the same reasons outlined above, likely be exacerbated in the field where even the average longevity of wild adults is estimated to be less than five days (33). Although such phenotypes would likely prove deleterious to those inheriting them in the field, the system as demonstrated here remains a ‘proof-of-principle’. For true ‘field-ready’ strains both a higher degree of penetrance and an earlier-acting phenotype would be required, especially for protection of cash-crops such as Brassicas where aesthetic damage by younger instars can prove economically damaging (2). Achieving this may prove as simple as generation of a larger number of tetO-AaHIT insertions in order to test for loci which allow earlier expression. For ease and economy of mass-rearing, it would also necessary to combine the bipartite tet-off components (e.g. *Hr5/ie1*-tTAV and tetO-*AaHIT*) into a single construct and genomic insertion site, although recent modelling has indicated that use of two-part systems can still function effectively at population regulation, at least when deleterious effects are limited to a single sex (34). Despite these caveats, our findings, alongside others, confirm the utility of the invertebrate-specific neurotoxic protein AaHIT as a means of effective insect control. More specifically, our results extend the potential of non cell-autonomous effectors as potential tools for GPM to the Lepidoptera, which include some of the world’s most damaging pest species.

## 5. Supporting Information

Supporting information is available at https://doi.org/10.6084/m9.figshare.12936839.v1. Supporting information includes video records of the ‘shaking’ phenotype observed in the *Hr5/ie1*>*AaHIT* and *Op/ie2*>*AaHIT* adults off-tet along with raw data used for all analyses.

## 6. Acknowledgements

T.H-S, V.C.N, R.C, E.L and L.AL were supported by European Union H2020 Grant nEUROSTRESSPEP (634361). T.H-S was additionally supported by a UK Biotechnology and Biological Sciences Research Council (BBSRC) Impact Acceleration Account grant (BB/S506680/1). PTL was funded through a Wellcome Trust Investigator Award [110117/Z/15/Z] made to LA. LA is supported by core funding from the UK Biotechnology and Biological Sciences Research Council (BBSRC) to The Pirbright Institute [BBS/E/I/00007033, BBS/E/I/00007038 and BBS/E/I/00007039]. X.X was supported by a CSC Scholarship from the Chinese government, and a PhD student exchange program from Fujian Agriculture and Forestry University (FAFU).

## References

1. Furlong MJ, Wright DJ, Dosdall LM. Diamondback moth ecology and management: problems, progress, and prospects. Annual Review of Entomology. 2013;58:517–41.

2. Zalucki MP, Shabbir A, Silva R, Adamson D, Liu S-S, Furlong MJ. Estimating the Economic Cost of One of the World’s Major Insect Pests, Plutella xylostella (Lepidoptera: Plutellidae): Just How Long Is a Piece of String? Journal of economic entomology. 2012;105(4):1115–29.

3. Talekar NS, Shelton AM. Biology, ecology, and management of the diamondback moth. Annual Review of Entomology. 1993;38:275–301.

4. Jin L, Walker AS, Fu G, Harvey-Samuel T, Dafa’alla T, Miles A, et al. Engineered female-specific lethality for control of pest Lepidoptera. ACS Synthetic Biology. 2013;2(3):160–6.

5. Alphey L. Re-engineering the sterile insect technique. Insect Biochemistry and Molecular Biology. 2002;32(10):1243–7.

6. Black WC, Alphey L, James AA. Why RIDL is not SIT. Trends Parasitol. 2011;27(8):362–70.

7. Harvey-Samuel T, Ant T, Gong H, Morrison NI, Alphey L. Population-level effects of fitness costs associated with repressible female-lethal transgene insertions in two pest insects. Evolutionary Applications. 2014;7(5):597–606.

8. Harvey-Samuel T, Morrison NI, Walker AS, Marubbi T, Yao J, Collins HL, et al. Pest control and resistance management through release of insects carrying a male-selecting transgene. BMC Biology. 2015;13, 49.

9. Zhou LQ, Alphey N, Walker AS, Travers LM, Morrison NI, Bonsall MB, et al. The application of self-limiting transgenic insects in managing resistance in experimental metapopulations. Journal of Applied Ecology. 2019;56(3):688–98.

10. Alphey N, Bonsall MB, Alphey L. Combining Pest Control and Resistance Management: Synergy of Engineered Insects With Bt Crops. Journal of economic entomology. 2009;102(2):717–32.

11. Alphey N, Coleman PG, Donnelly CA, Alphey L. Managing insecticide resistance by mass release of engineered insects. Journal of economic entomology. 2007;100(5):1642–9.

12. Bolton M, Collins HL, Chapman T, Morrison NI, Long SJ, Linn CE, et al. Response to a Synthetic Pheromone Source by OX4319L, a Self-Limiting Diamondback Moth (Lepidoptera: Plutellidae) Strain, and Field Dispersal Characteristics of its Progenitor Strain. Journal of Economic Entomology. 2019;112(4):1546–51.

13. Shelton AM, Long SJ, Walker AS, Bolton M, Collins HL, Revuelta L, et al. First Field Release of a Genetically Engineered, Self-Limiting Agricultural Pest Insect: Evaluating Its Potential for Future Crop Protection. Front Bioeng Biotech. 2020;7.

14. Phuc HK, Andreasen MH, Burton RS, Vass C, Epton MJ, Pape G, et al. Late-acting dominant lethal genetic systems and mosquito control. BMC Biology. 2007;5:11-.

15. Gossen M, Bujard H. Tight Control of Gene-Expression in Mammalian-Cells by Tetracycline-Responsive Promoters. Proceedings of the National Academy of Sciences of the United States of America. 1992;89(12):5547–51.

16. Bryk J, Reeves RG, Reed FA, Denton JA. Transcriptional effects of a positive feedback circuit in Drosophila melanogaster. Bmc Genomics. 2017;18.

17. Fu GL, Lees RS, Nimmo D, Aw D, Jin L, Gray P, et al. Female-specific flightless phenotype for mosquito control. Proceedings of the National Academy of Sciences of the United States of America. 2010;107(10):4550–4.

18. Labbe GMC, Scaife S, Morgan SA, Curtis ZH, Alphey L. Female-Specific Flightless (fsRIDL) Phenotype for Control of Aedes albopictus. Plos Neglected Tropical Diseases. 2012;6(7).

19. Marinotti O, Jasinskiene N, Fazekas A, Scaife S, Fu G, Mattingly ST, et al. Development of a population suppression strain of the human malaria vector mosquito, Anopheles stephensi. Malaria Journal. 2013;12:142.

20. Haghighat-Khah RE, Harvey-Samuel T, Basu S, StJohn O, Scaife S, Verkuijl S, et al. Engineered action at a distance: Blood-meal-inducible paralysis in Aedes aegypti. Plos Neglected Tropical Diseases. 2019;13(9).

21. Deng SQ, Chen JT, Li WW, Chen M, Peng HJ. Application of the Scorpion Neurotoxin AaIT against Insect Pests. Int J Mol Sci. 2019;20(14).

22. Martins S, Naish N, Walker AS, Morrison NI, Scaife S, Fu G, et al. Germline transformation of the diamondback moth, Plutella xylostella L., using the piggyBac transposable element. Insect Molecular Biology. 2012;21(4):414–21.

23. Fu W, Xie W, Zhang Z, Wang SL, Wu QJ, Liu Y, et al. Exploring Valid Reference Genes for Quantitative Real-time PCR Analysis in Plutella xylostella (Lepidoptera: Plutellidae). International Journal of Biological Sciences. 2013;9(8):792–802.

24. Perkins JR, Dawes JM, McMahon SB, Bennett DLH, Orengo C, Kohl M. ReadqPCR and NormqPCR: R packages for the reading, quality checking and normalisation of RT-qPCR quantification cycle (Cq) data. Bmc Genomics. 2012;13.

25. Hellemans J, Mortier G, De Paepe A, Speleman F, Vandesompele J. qBase relative quantification framework and software for management and automated analysis of real-time quantitative PCR data. Genome Biol. 2007;8(2):R19.

26. Therneau MT, Grambsch PM. Modeling Survival Data: Extending the Cox Model. New York: Springer; 2000.

27. Kassambara A, Kosinski M, Biecek P. survminer: Drawing Survival Curves using ggplot2 R package version 0.4.6.: https://CRAN.R-project.org/package=survminer; 2019

28. Hothorn T, Bretz F, Westfall P. Simultaneous inference in general parametric models. Biom J. 2008;50(3):346–63.

29. Wickham H, Averick M, Bryan J, Chang W, D’agostino McGowan L, Francois R, et al. Welcome to the tidyverse. Journal of Open Source Software. 4(43).

30. Kokoza V, Ahmed A, Cho WL, Jasinskiene N, James AA, Raikhel A. Engineering blood meal-activated systemic immunity in the yellow fever mosquito, Aedes aegypti. Proceedings of the National Academy of Sciences of the United States of America. 2000;97(16):9144–9.

31. Kokoza VA, Raikhel AS. Targeted gene expression in the transgenic Aedes aegypti using the binary Gal4-UAS system. Insect Biochemistry and Molecular Biology. 2011;41(8):637–44.

32. Romans P, Tu ZJ, Ke ZX, Hagedorn HH. Analysis of a Vitellogenin Gene of the Mosquito, Aedes-Aegypti and Comparisons to Vitellogenins from Other Organisms. Insect Biochemistry and Molecular Biology. 1995;25(8):939–58.

33. Furlong MJ, Pell JK, Pek Choo O, Abdul Rahman S. Field and laboratory evaluation of a sex pheromone trap for the autodissemination of the fungal entomopathogen Zoophthora radicans (Entomophthorales) by the diamondback moth, Plutella xylostella (Lepidoptera: Yponomeutidae). Bulletin of entomological research. 1995;85(03):331–7.

34. Vella MR, Gould F, Lloyd AL. Mathematical modeling of genetic pest management through female lethality with independently segregating alleles. BioRxiv. 2020.

